# Structure of phage λ Redβ_177_ annealase shows how it anneals DNA strands during single-strand annealing homologous DNA recombination

**DOI:** 10.1101/2022.04.09.487726

**Authors:** Timothy Newing, Jodi L. Brewster, Haibo Yu, Nikolas P. Johnston, Lucy J. Fitschen, Gökhan Tolun

## Abstract

The bacteriophage λ red recombination system catalyzes the single-strand annealing homologous DNA recombination reaction, in which Redβ annealase protein plays a critical role. Using cryogenic electron microscopy, we were able to determine a structure of a C-terminally truncated Redβ with the residues 1-177 bound to two complementary 27mer oligonucleotides forming an annealing intermediate, to a final resolution of 3.3 Å. This structure validates and rationalizes decades of experimental observations on the biochemistry of Redβ. Definition of the interaction surfaces between subunits explains not only the DNA binding properties of Redβ, but also its propensity to oligomerize into long helical filaments, which are also formed by almost all annealases and are known to be functionally important. In addition, this annealing intermediate structure provides a detailed picture of the hydrogen bonding network that positions the DNA strands in a planar orientation to facilitate base pairing. Residues 133-138, which are missing from our structure, form a flexible loop. Molecular dynamics simulations were used to model the range of motion of the flexible loop, which suggested that it has a crucial role in keeping the DNA strands in the DNA-binding groove long enough to allow homology searching. The combination of structural and computational observations has allowed us to propose a detailed mechanism for the action of Redβ. More than half a century after its discovery, our work shines a light not only on the structure and mechanisms of Redβ, but also of other proteins within the annealase superfamilies.

**Significance Statement:** Single-strand annealing homologous DNA recombination is a process that is conserved throughout evolution from bacteriophages to humans, highlighting its importance and indispensability. It is a process that repairs double-stranded DNA breaks and is particularly vital in dsDNA viruses. The Redβ protein from the bacteriophage lambda is the archetypal annealase protein, forming the basis of our knowledge about this class of proteins. Along with the exonuclease λExo, these two proteins not only form the model system for single-strand annealing homologous recombination, but are also used in thousands of laboratories worldwide for performing genetic manipulations. After its discovery in 1966, we report the first structure of the DNA-binding and oligomerization domain of Redβ, providing details about the mechanism of homologous DNA annealing.

## Introduction

Damage to DNA in the form of double-stranded breaks may occur due to the action of endonucleases, at chromosomal crossovers, at stalled or reversed replication forks, or due to environmental agents such as chemicals and radiation (*1*). Single-strand annealing homologous recombination (SSA) enables dsDNA breaks to be repaired, which is vital for maintaining genomic fidelity. Simultaneously, it is an inherently mutagenic process and provides an important source of genetic diversity (*2*). SSA is evolutionarily conserved and it is present in many organisms, from simple bacteriophages to humans (*3*).

The Red system from bacteriophage lambda (phage λ) is comprised of the genes *exo, bet* and *gam*, which encode for the proteins λ exonuclease, Redβ and γ-protein, respectively. The Red system proteins are capable of repairing dsDNA breaks by SSA. λ exonuclease (λExo [originally, Redα]) is a 25.9 kDa protein that forms a toroidal homotrimer (*4*), which binds to dsDNA ends. It resects one of the strands in the 5’ to 3’ direction, converting the other stand to a 3’ ssDNA overhang (*5-7*). Redβ (a.k.a. Beta), a 29.7 kDa protein binds to and protects this exposed nascent ssDNA, catalyzing its annealing to a homologous DNA molecule (*8*). λExo and Redβ have been shown to interact in a 1:1 monomer:monomer stoichiometry to form an **<u>E</u>**xonuclease **<u>A</u>**nnealase **<u>T</u>**wo-component **<u>R</u>**ecombinase (EATR) complex (a.k.a. SynExo) (*9-11*). The third protein, γ (Gam), though not physically associated with EATR, binds to and inhibits host *Escherichia coli* RecBCD nuclease (*12, 13*).

The phage λ Red system is the archetypal EATR complex and has been studied extensively over the last 50 years as a model system for SSA. It has also emerged as a tool for genetic manipulation, termed recombineering (*14, 15*). There are currently two proposed models for how SSA may occur, which have been recently reviewed by us (*16*) and others (*1, 17*). In brief, variable processing of dsDNA ends by λExo leads to two different scenarios. These scenarios describe either annealing between two partially processed dsDNA ends for repair independent of DNA replication; or complete removal of one of the DNA strands leaving a ssDNA intermediate that is incorporated into the lagging strand at a DNA replication fork. These models were initially proposed by Lin *et al*. (*18*) and Mosberg *et al*. (*19*), respectively. In both instances, Redβ is proposed to mediate homology searching and the annealing of homologous sequences. A version of the latter model forms the basis of another genetic tool, **<u>M</u>**ultiplex **<u>A</u>**utonomous **<u>G</u>**enome

**<u>E</u>**ngineering (MAGE) (*20*), for the simultaneous targeted mutagenesis of numerous genes. However, many questions related to the process of SSA remain unanswered, particularly regarding the detailed molecular mechanisms underlying the catalysis of homology searching and annealing by Redβ.

Functional homologs of Redβ appear to be almost ubiquitous, with an increasing number of representatives being discovered across viral, bacterial, and eukaryotic domains. Despite this, few of the annealase proteins have received even rudimentary characterization. Therefore, there is a paucity of information regarding the biochemical, structural and functional attributes of annealases. However, the available data on Redβ and other annealases such as ERF (phage P22), ORF6 (Kaposi’s Sarcoma Herpes Virus), ICP8 (Herpes Simplex Virus-1), RecT (*E. coli*), human Rad52 and its eukaryotic homologs suggest some commonalities. Annealases are ATP-independent and do not catalyze strand invasion of duplex DNA; thus, they are functionally distinct from the well-known and studied ATPase recombination proteins, such as RecA from *E. coli* and eukaryotic Rad51 (*21, 22*). However, in a manner reminiscent of *E. coli* SSB, annealases bind to ssDNA, holding it in an extended conformation, removing secondary structures and protecting it from degradation (*23*). The DNA binding properties of Redβ differ substantially from those of SSB. Redβ binds only weakly to ssDNA, but in the presence of magnesium, it binds stably and with high affinity to a nascent dsDNA, the product of annealing, such as that formed upon the sequential addition of two complementary ssDNA oligos (*24*). Redβ does not bind at all to preformed duplex DNA (*24*).

Redβ is known to form ring-like and filamentous oligomeric superstructures that are proposed to be functionally important. Electron and atomic force microscopy studies have provided images and low-resolution structural information on the oligomeric complexes of Redβ. In the absence of DNA, Redβ forms an abundance of small rings or split-lock washer conformers, each with an average of 11-12 subunits (*25, 26*). However, in the presence of ssDNA, Redβ forms a heterogeneous mix of 11-19 subunit rings, distorted and part rings, and short left-handed helical filaments (*11, 25, 26*). In the presence of heat-denatured complementary DNA, the larger rings and filament superstructures are stabilized and are proposed to represent an annealing intermediate (*24-26*). The oligomeric heterogeneity of Redβ was recently expanded upon through a detailed, native mass spectrometry analysis (*27*). Above a protein concentration of 1 μM, numerous oligomeric states from 5 to 14 subunits were detected for both Redβ alone and in the presence of short ssDNA and complementary oligonucleotides. Increasing the Redβ concentration up to 30 μM (still determined to be within the physiologically relevant range) promoted larger structures, as did an increase in the length of DNA used (*27*).

Like Redβ, other annealases have a propensity to form similar ring-like quaternary structures and helical filament superstructures, which appear to suggest a common mode of action. In addition, rings are often seen at the termini of long helical filaments (*25, 28*), giving rise to the hypothesis that the ring and helical forms interconvert (*25, 29*). Many such oligomeric complexes have been observed by electron microscopy, including Orf6 (*30*), Rad52 (*22, 31-34*) and ICP8 (*28, 35-38*). Low-resolution reconstructions of ICP8, both in the form of binary nonameric rings (in the presence of short ssDNA) (*29*) and bipolar filaments (*39*) have been generated(*29, 39*). The bipolar filaments form in both the absence of DNA (*36, 38, 39*) and the presence of heat-denatured DNA (*28, 37*), and likely represent an intertwining of single filaments seen in the presence of ssDNA (*35*). The ring and bipolar filaments have also been proposed as interconverting annealing intermediates (*28, 29, 39*). The fitting of an available crystal structure of the ICP8 monomer, with a 60 residue C-terminal truncation (*40*), into the reconstructions has provided information on intermolecular contacts involved in oligomerization.

Though they share functional and oligomerization properties, the annealases of bacterial and phage origin appear to form a group distinct from, and likely evolutionarily unrelated to, the eukaryotic-host viral annealases, such as ICP8, and homologs BALF2 and ORF6. The latter are much larger at ∼130 kDa, compared to 29.7 kDa for Redβ. This difference in size may reflect a diversity of function and a greater number of interacting partners. In addition to involvement in SSA, ICP8 is proposed to have multiple roles in DNA replication, replication compartment formation and gene expression (*41, 42*). Phylogenetic analyses of key annealases, including human, bacterial and phage derived proteins, appear to resolve at least three evolutionary lineages, each forming a superfamily: Gp-like, Rad51-like and Rad52-like (*43*). Later publications further divided these lineages into five distinct families named for their respective flagship members Sak3, Sak4, Erf, Rad52 and RecT/Redβ (*44*). The only near-atomic resolution structural data of an oligomeric complex available amongst these families is of the undecameric rings of the Rad52 N-terminal domain (residues 1-212) (*32, 45, 46*). This data revealed an inner groove and an outer groove on the ring surface, where ssDNA was bound. The two grooves were suggested to represent two different modes of DNA binding and reaction intermediates. The deep inner groove, predicted to be the site of annealing, is lined with positively charged arginine and lysine residues that interact with the phosphate backbone of ssDNA, positioning the nucleotides outwards for complementary base pairing.

The architecture of annealase proteins can be crudely separated into two functionally distinct domains. The N-terminal domain (NTD) mediates oligomerization and contains the catalytic functionality. Residues 1-177 of Redβ (Redβ_177_) are sufficient for binding to ssDNA (*47*) and catalyzing the annealing of complementary strands *in vitro* (*11, 48*).

Although there are some suggestions that the C-terminal domain (CTD) may modulate the strength of DNA interactions, it does not appear to be required (*49*). The much smaller CTDs are implicated in mediating protein-protein interactions. Residues 178-261 of Redβ interact with λExo (*11*) and potentially other host proteins such as SSB (*49*).

The NTD and CTD of annealases are flexible relative to one another, which presents major challenges for structure determination. The current available crystal structures of both ICP8 (*40*) and Rad52 (*32, 45, 46*) are C-terminally truncated. However, while these crystal structures provide tantalizing insights into the function of annealase proteins within that family, they are of limited use for broader extrapolation. The amino acid sequence similarity between annealases is very low, often less than 15-20% (*43*), making sequence alignments less reliable and impractical for the purpose of generating homology models for annealases in different families.

An X-ray crystal structure of the C-terminal domain of Redβ (residues 183-261) in complex with the λExo toroid showed a compact 3-helix bundle (*50*). However, there is no representative structure of the NTD, the main domain that carries out the primary functions of DNA binding, annealing and homo-oligomerization required for SSA. As Redβ is undoubtedly the archetype annealase for its family, the absence of structural information has, until now, been a major barrier in our understanding of not only bacteriophage annealases, but also how SSA functions.

Here we present a structure of the helical annealing intermediate of Redβ_177_, in complex with complementary oligonucleotides, determined by cryo-electron microscopy to 3.3 Å. Our structure allows the first detailed look at both the nucleoprotein interactions and the interprotein contacts dictating the formation of the helical superstructure it forms. This structure is the first that demonstrates the mechanism of action for Redβ_177_, while also having the potential to be used as a template for homology models for other members of this protein superfamily. Combined with CTD crystal structure and the structure prediction we made using AlphaFold v2.0 (*51*) we finally present a complete picture of the Redβ structure more than half a century after its discovery.

## Results

We present the first-ever structure of the DNA-binding and oligomerization domain of Redβ annealase from phage λ. This structure, resolved to 3.3 Å, was obtained by cryo-EM via helical reconstruction, using a C-terminally truncated Redβ_177_ bound to two 27mer complementary oligonucleotides.

### Reconstructed helical cryo-EM density map

The Redβ_177_ was expressed in *E. coli* and purified by immobilized metal ion affinity chromatography followed by gel filtration chromatography (Fig. S1). The purified protein was incubated with oligonucleotides before being plunge frozen for cryo-EM. We collected 4,701 micrographs, from which 259,649 helical particles were identified using template-based helical filament tracing and subsequent particle curation in cryoSPARC (*52*). The micrographs contained helices in a variety of conformations, but most were uniform, compact “telephone cord-like” solenoids, though some sections were less condensed (Fig. 1A). The 2D class averages generated during the reconstruction (Fig. 1B and Fig. S2A), together with their Fourier transformation (Fig. 1C), clearly show the features of the helical structures formed by Redβ_177_. Two subpopulations of particles, representing different helical symmetries, were identified in 2D classification – a 1-start and a 2-start helical assembly, consisting of 26,525 and 77,799 particles, respectively (Fig. S2). Previous work has established that Redβ forms left-handed helices (*25*), therefore left-handed helical symmetry was imposed during further processing.

**Fig. 1.**
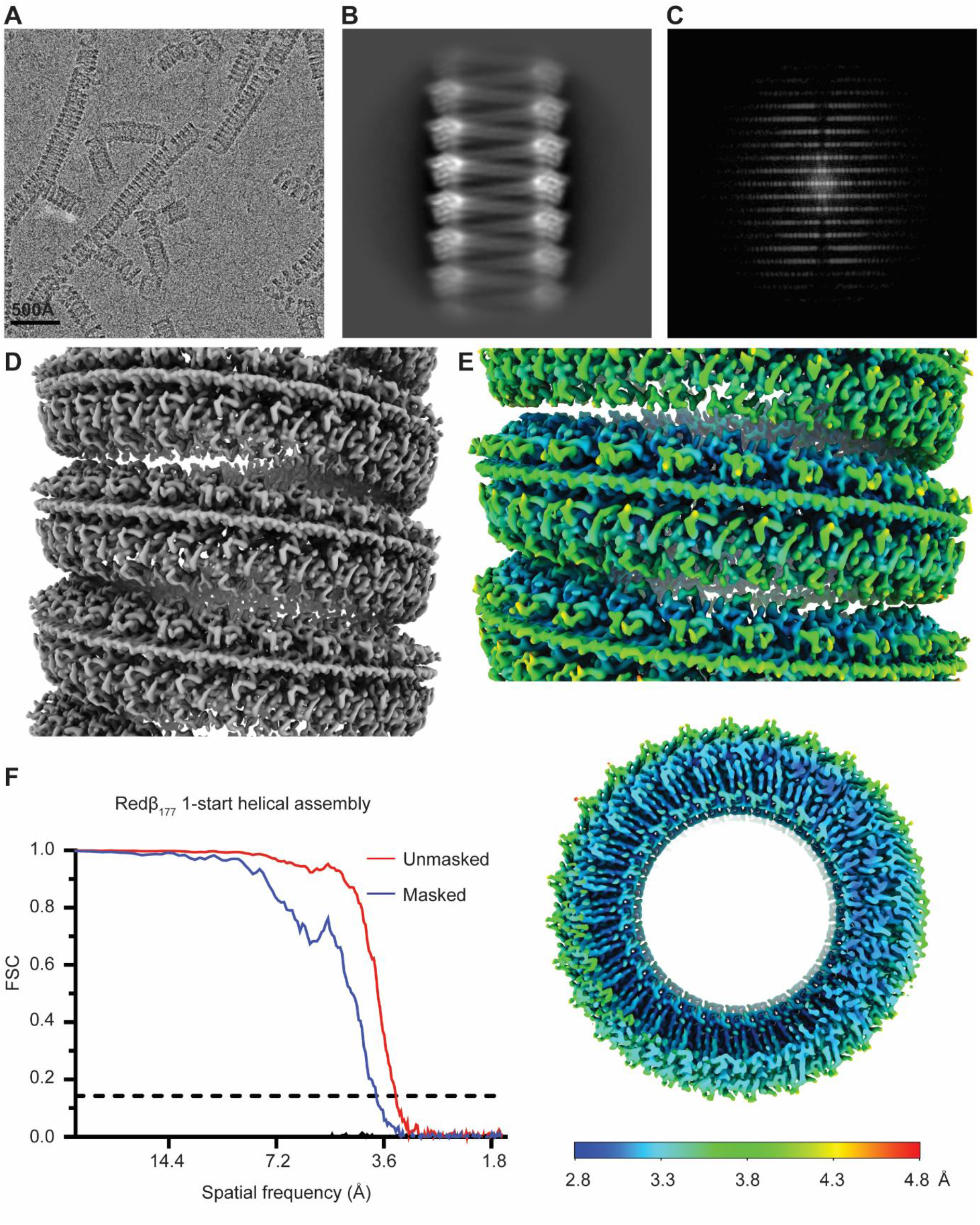
Cryo-EM structure of Redβ_177_ 1-start helical assembly, resolved to 3.3 Å. (**A**) A representative electron microscopy micrograph of the Redβ_177_ helices. (**B**) A representative 2D class average of the helical filaments. (**C**) Fourier transform of a 2D class average. (**D**) Cryo-EM electron density map of the 1-start helical assembly resolved to 3.3 Å. (**E**) Local resolution of the structure visible from both a side and a top view. (**F**) Plot showing the Fourier shell correlation (FSC) versus spatial frequency, for both the masked (blue) and unmasked (red) maps. The resolution of the structure was assessed using the point where the FSC curve crosses a correlation value of 0.143.

Helical reconstruction and subsequent symmetry determination of the 1-start helical population identified a twist of -12.947 degrees and a rise of 2.078 Å. Following refinement of the local motion and CTF, symmetrized, non-uniform helical refinement of the 1-start particles resulted in a map with an estimated final resolution of 3.3 Å (Fig. 1D-F), according to the gold-standard FSC = 0.143 (GSFSC) criteria (*53*). However, the local resolution of the map, as determined by cryoSPARC, ranges from 2.8 Å to 4.8 Å (Fig. 1E). The overall image processing workflow can be seen in Fig. S3 and structure statistics can be found in Table S1. A single turn of the 1-start helix contains 27.8 asymmetric units, each corresponding to a Redβ_177_ monomer (Fig. 2A). The density of double-stranded DNA, which represents an average of possible DNA sequences bound to a monomer of Redβ_177_, can be seen wrapping around the circumference of the protein helix (Fig. 1D, 1E, 2A). The presence of double-stranded DNA in the map indicates that we have captured an annealing intermediate of the SSA homologous recombination reaction. The attempt to reconstruct the structure of the 2-start helical complex did not yield a reliable map, likely due to the presence of irresolvable continuous variation in the helical symmetry parameters. Therefore, we are presenting only the 2D class averages that indicate a helical assembly composed of two apparently identical Redβ_177_ filaments (Fig. S2B).

**Fig. 2.**
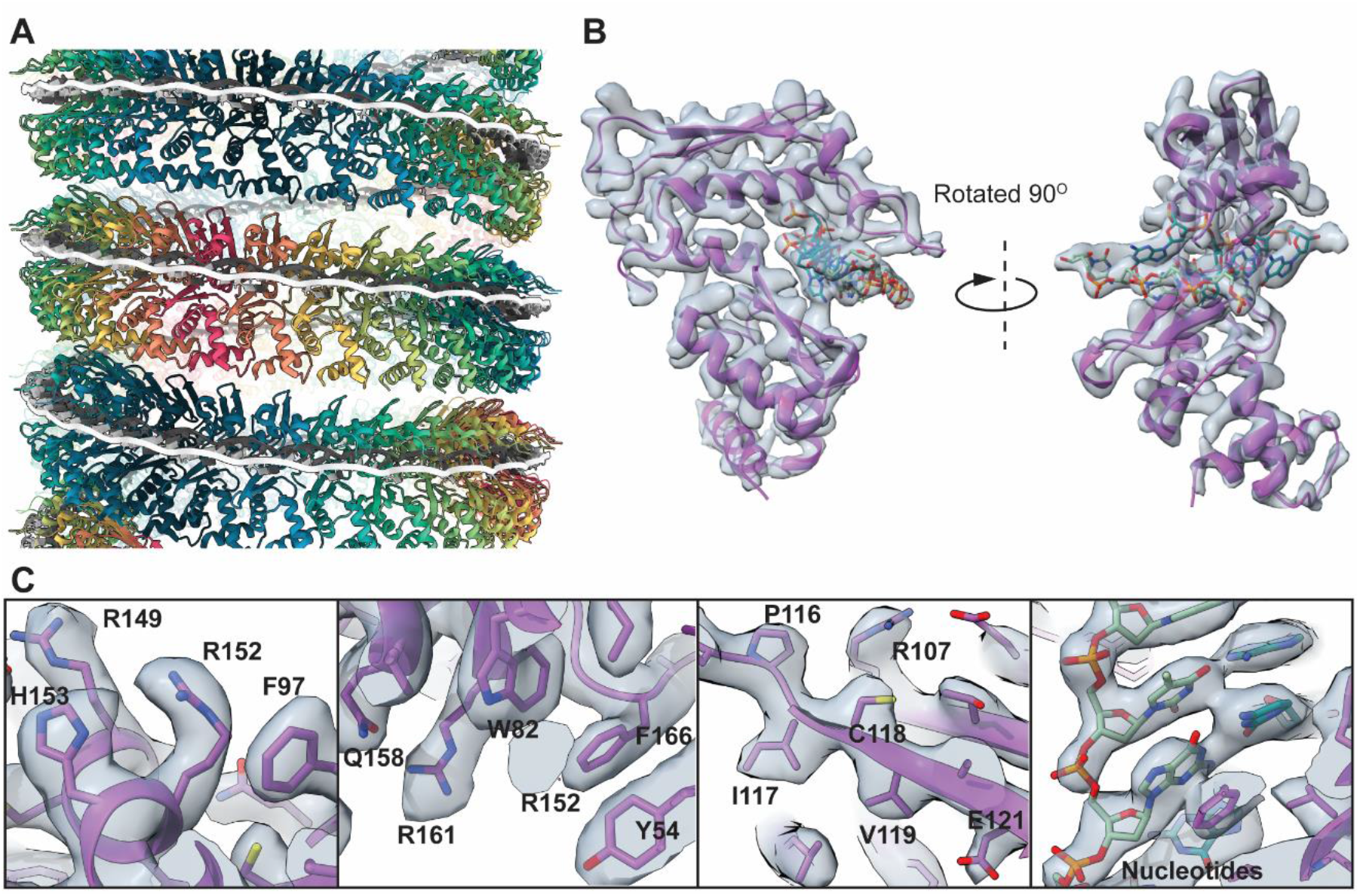
Redβ_177_ and DNA structure. (**A**) The ribbon structure of the Redβ_177_ helix showing the interaction between protein and DNA (black and white) in a planar conformation. (**B**) The electron map of a single Redβ_177_ monomer (grey) bound to DNA, with the ribbon structure within it (purple). Close up views of this monomer (**C**) demonstrates the fit of amino acids and nucleotides within the density map.

### Atomic model confirms Redβ_177_ is structurally similar to human Rad52

A total of 161 residues of Redβ_177_ were built *de novo* into the density (Fig. 2B). This encompassed residues 3-169, except for a flexible loop region (residues 133-138) which could not be accurately modelled (see below). The density for residues 1-2 and 170-177 was also insufficient quality to allow modelling. However, the quality of the map allowed for the accurate positioning of the sidechains of majority of the remaining residues and of the oligonucleotides (Fig. 2C). The coordinates for the atomic model have been deposited into the PDB (7UJL) and the map into EMDB (EMD-26566). In general, the Redβ N-terminal domain consists of a mixed αβ fold consisting of five α-helices, a 3_10_ helix and five β-strands, which form two separate β-sheet elements (Fig. 3A). The structure contains a recognizable βββα core motif (β3-5 and α5), which is a small OB-fold, present in DNA-binding proteins (*54*). This motif is also present in the crystal structure of the N-terminal domain of Rad52 (Fig. 3B), where it contributes to the formation of the deep-groove of the inner DNA-binding pocket (*46*). However, in Redβ_177_ the β-strands are much shorter, 5-6 amino acids in length, relative to those of the Rad52 βββα motif (10-14 amino acids in length). As a result, the outward-facing DNA-binding groove on the surface of Redβ_177_ is much shallower and more exposed than that of Rad52 (Fig 3A, 3B). A structural similarity search with Redβ_177_, using the DALI server (*55*) identified Rad52 as the most similar structural fold. This is consistent with prior work placing Redβ and many other bacterial annealases in a superfamily that is proposed to share a common fold (*43*).

**Fig. 3.**
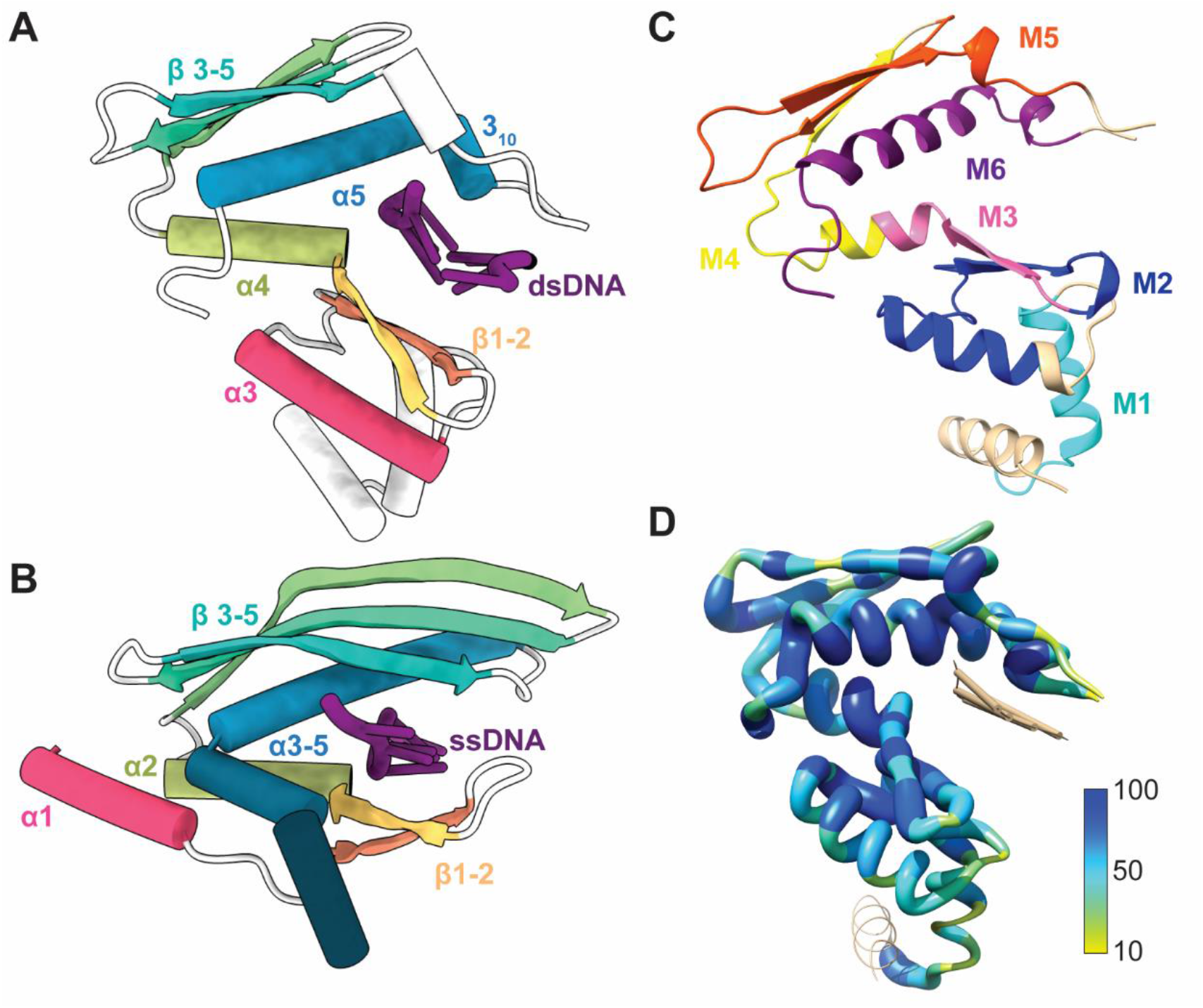
Conserved annealase structural motifs. Cartoon models comparing the secondary structures of Redβ_177_ (**A**) and Rad52 (**B**). Color denotes conserved secondary structures between the proteins, while white represents regions that are not conserved between the structures and DNA is shown in purple. (**C**) Demonstrates the ribbon structure of the Redβ177 monomer, with colors indicating conserved motifs 1-6 that were identified by the MSA (Fig. S4). Beige indicates no conservation. (**D**) Amino acid sequence conservation of Redβ_177_ compared to other annealases. The extent of sequence conservation (%) is denoted by thickness and color.

### Multiple amino acid sequence alignment identifies conserved motifs

Using the Redβ protein sequence (UniProtKB: P03698) as a reference, we constructed a multiple sequence alignment (MSA) based on the top 1000 hits from a BLAST search of the UniProt Reference Cluster (UniRef) 90% database. Following refinement, our MSA consists of 425 reference cluster sequences and is 411 residues in length (Fig. S4). From this alignment, we identified eight conserved motifs based on >50% consensus across the MSA (Fig. S4). Six of these motifs were able to be mapped to the structure of Redβ_177_ (Fig. 3C and Fig. S5), the other two motifs are located in the C-terminal domain. The amino acid sequence conservation was also mapped onto the structure of Redβ177 (Fig. 3D). This showed that the most highly conserved residues, which are in motifs 2, 3 and 6, form the DNA-binding pocket. Two residues in particular, F66 and W143, are almost universally conserved across the sequences in the database.

### Oligomerization

The model of Redβ_177_ was reconstructed as a compact, solenoid, helical filament with dsDNA wound around the outside. This superstructure is stabilized through substantial electrostatic contacts between adjacent monomers in the complex (Fig. 4A, 4B, Fig. S6). Much of the large interaction surface area is comprised of patches of positive and negative charge, which are mirrored on the adjacent monomer face (Fig. 4B). In particular, a cluster of strong electrostatic interactions formed by E125, E121, K148’, and R149’; R161 and D80’; D68, K61’ and K36’, may help to stabilize the area around the DNA-binding pocket (Fig. 4B). Some of these residues such as K36, K61, and R161 are also involved in DNA-binding. Therefore, oligomerization of adjacent units may also be indirectly facilitated by the binding of the DNA substrate, as described below.

**Fig. 4.**
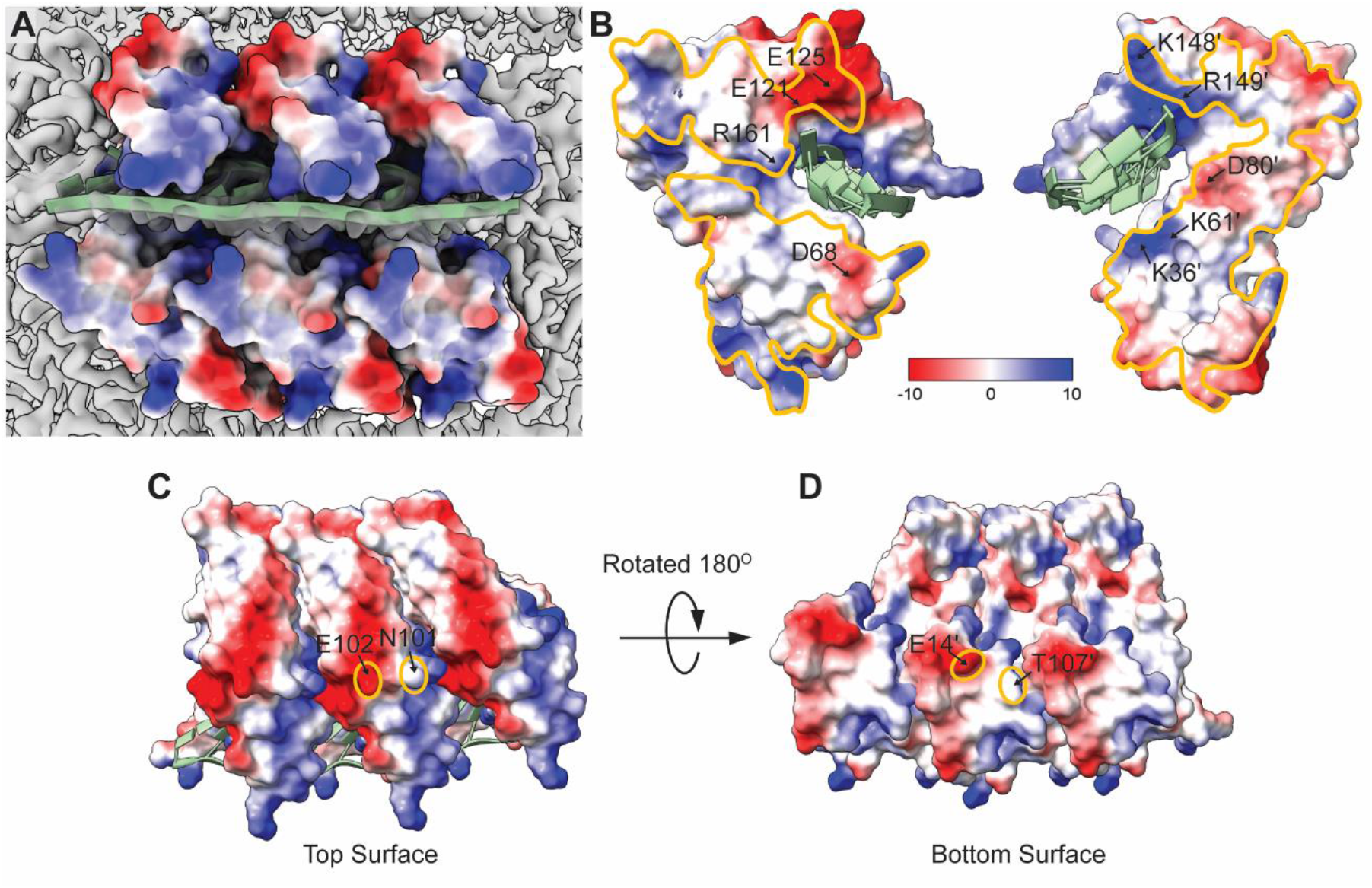
Electrostatic surface representation of Redβ_177_. The surface is colored by charge with positive (red), negative (blue) and electroneutral (white). (**A**) Electrostatic representation of three monomers Redβ_177_ bound to dsDNA (pale green). (**B**) The electrostatic interactions between two adjacent monomers within the helix. The orange line highlights the areas where the monomers interact (**C**-**D**). Interactions between the top (**C**) and bottom (**D**) surfaces of the Redβ_177_ monomers within the helix. The orange ovals highlight the interacting residues.

The interacting bottom-to-top monomer surfaces contribute relatively little to the stabilization of the helix (Fig 4 C, D). There are only two points of contact, E102 and N101; and T107’ and E14’, between the top and bottom surfaces, respectively. Therefore, the side-by-side contacts are the important determinants of helical characteristics such as the pitch and rise. Likewise, the abundance of electrostatic interactions between adjacent monomers drives elongation of the helical fragment.

### DNA binding properties of Redβ_177_ and the mechanism for the catalysis of DNA annealing

In order to capture an annealing intermediate structure, which would shed light on the mechanism of SSA, our sample was prepared for cryo-EM by sequentially incubating Redβ_177_ with complementary ssDNA oligonucleotides. This approach has been previously used for Redβ in *in vitro* experiments to form annealing intermediates (*25-27*) and it resulted in clear density for a double-stranded DNA annealing intermediate in our structure. The outward-facing DNA-binding groove accommodates both strands of the annealed dsDNA product, with an observed stoichiometry of four base pairs per Redβ_177_ monomer (Fig. 5A). However, adjacent monomers bind the template strand in an overlapping manner, such that a single monomer makes contact with nucleotides involved in forming 6 base pairs. The DNA binding groove is lined with positively charged residues (Fig. 4B), which accommodate the negatively charged phosphate backbone of the DNA, through a combination of electrostatic interactions and hydrogen bonds (Fig. 5B).

**Fig. 5.**
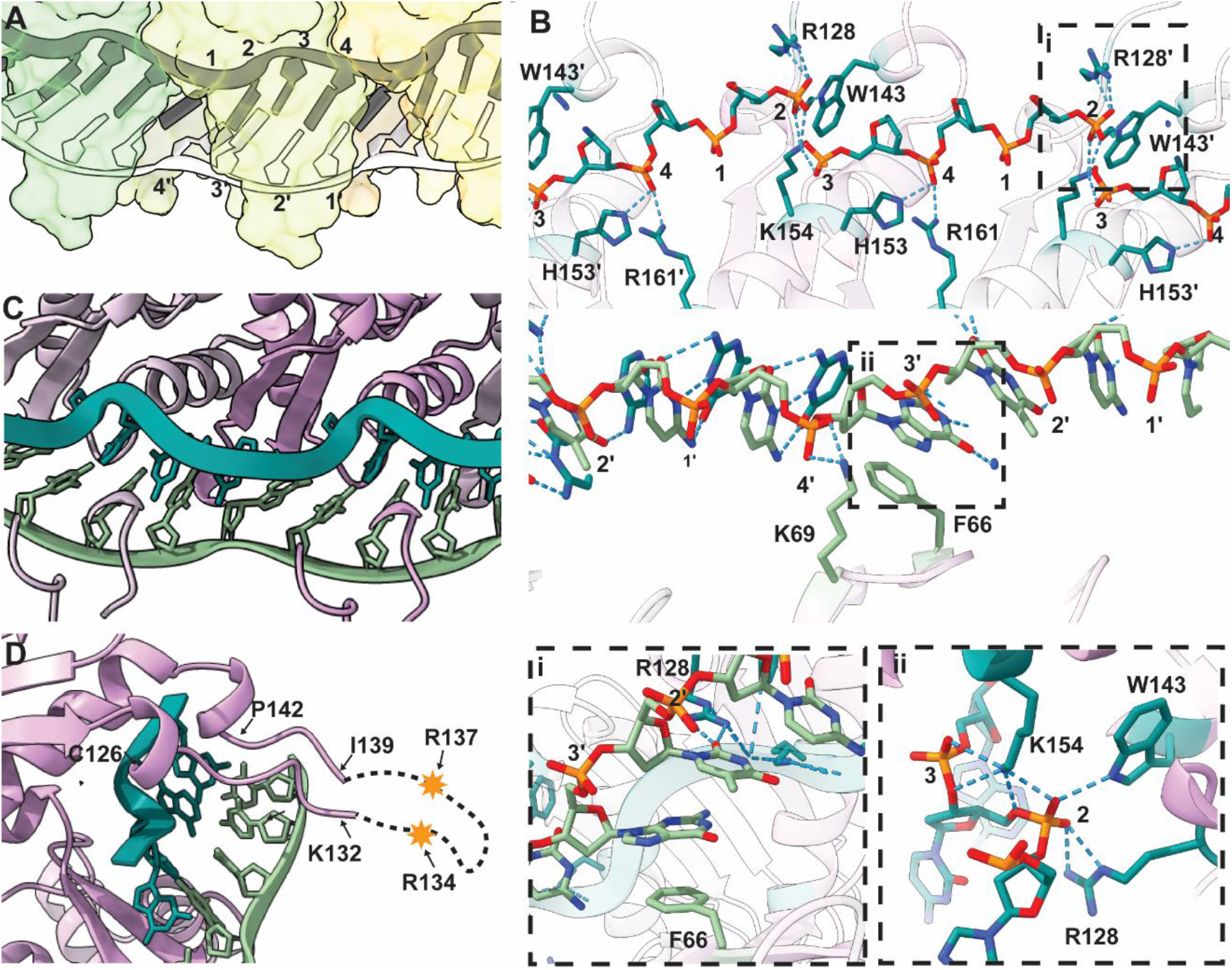
DNA binding to Redβ_177_. dsDNA binds to Redβ_177_ with a site size of 4, as numbered in (**A**), at the position of the phosphate of each nucleotide. (**B**) Hydrogen bonding between Redβ_177_, the template (top panel) and incoming (bottom panel) DNA strands. Close-up views of the π-stacking interaction between F66 and a nucleotide (**i**) and the H-bonding network stabilizing the kink in the phosphate backbone (**ii**) are shown. Ribbon diagrams of DNA (green) binding to three Redβ_177_ monomers (purple) clearly show the kink in the phosphate backbone of the template strand (**C**) as well as the finger loop (**D**), highlighting the residues which are missing from the structure. A prime (’) symbol denotes numbering in an adjacent molecule.

A total of nine hydrogen bonds between the phosphate backbone and the side chains of R128, W143, H153, K154 and R161 secure the inner, template ssDNA strand in the DNA-binding groove of Redβ_177_ (Fig. 5B, top panel). These bonds result in the nucleotides adopting a planar orientation and facilitate solvent exposure to provide a template for base-pairing to incoming homologous ssDNA. In contrast, the incoming ssDNA strand is only stabilized by two hydrogen bonds to the phosphate backbone through K69 (Fig. 5B bottom panel). In addition, the incoming strand is stabilized in a planar orientation through a π-stacking interaction between the phenyl ring of F66 and the purine or pyrimidine ring of the position 3’ nucleotide and two hydrogen bonds between R128 and the adjacent nucleotide at position 2’ (Fig 5B panel i). These interactions may tether and orient the incoming ssDNA to the annealing complex, whilst allowing it to slide against the template strand until regions of base pairing stabilize the duplex. The annealed duplex DNA is also stabilized in an extended, planar conformation in the DNA-binding groove (Fig. 5), which is atypical of B-form helical dsDNA conformations often found in solution (*56*). It is likely that upon annealing, the planar conformation is maintained through an induced distortion in the phosphate backbone of the inner (template) DNA strand, which is stabilized by a network of seven hydrogen bonds with R128, W143 and K154 (Fig 5B panel ii). This kink (Fig. 5C) is necessitated due to the shorter helical radius, and thus shorter path of the inner DNA strand, relative to the outer strand.

Residues 133-138 could not be modelled in our structure. This region forms part of a larger, positively charged, loop (residues 126-142) that forms the roof of the DNA-binding groove (Fig. 5D). It is likely that residues 133-138 that form the tip of the loop are highly flexible, the motion of which (i.e., conformational variability) precluded the formation of structured density for this region during reconstruction. To see how this loop may be moving and functioning, we ran molecular dynamics (MD) simulations using a trimeric assembly of apo-protein (Redβ_177_ only), Redβ_177_ bound either to template ssDNA or to both DNA strands. Molecular dynamics simulations were run both with weak harmonic restraints applied on the two outside monomer and DNA backbone atoms (Fig. 6) and also without any restraints, as a control (Fig. S7). Weak harmonic restraints help to maintain the oligomeric Redβ structure (modelled by a trimeric structure) and the intermediate DNA structure. Each system was simulated for 1 μs during which the RMSD of the protein backbone positions and RMSF of the Cα atom of the central trimeric subunit were calculated (Fig. S7D-G).

**Fig. 6.**
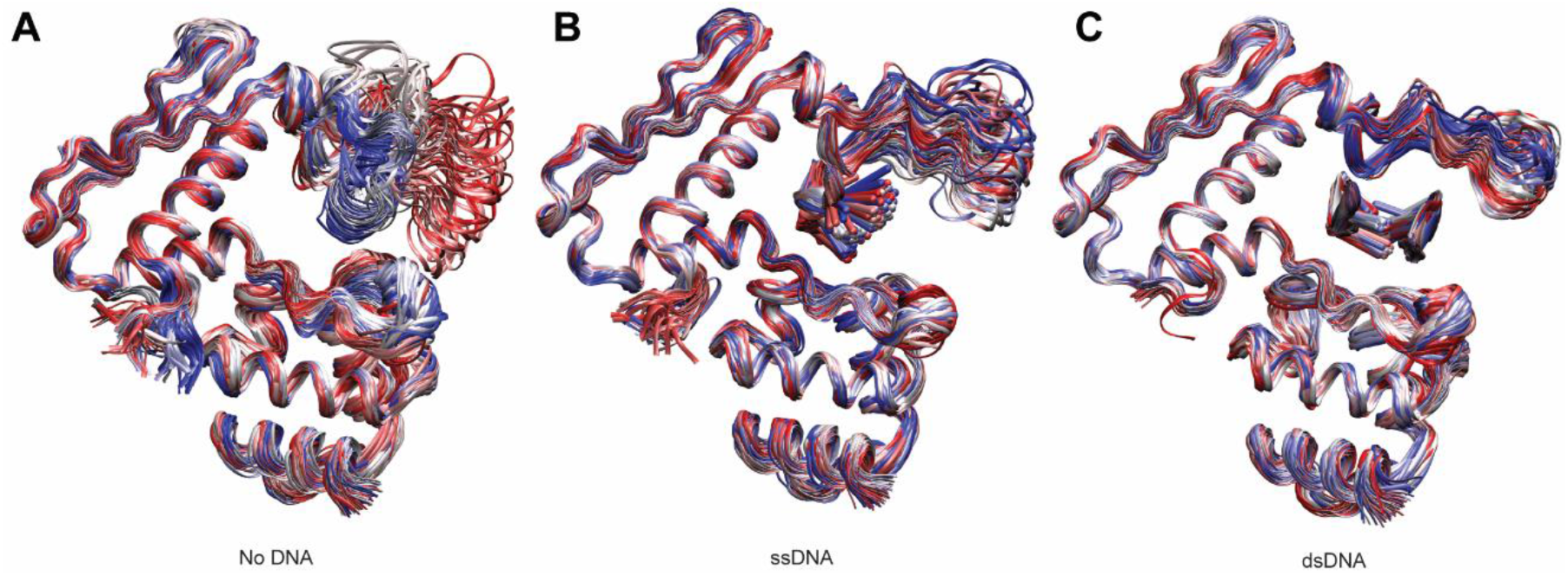
Molecular Dynamics simulations of Redβ_177_. The sampled ensemble of apo (**A**), ssDNA-bound (**B**) and dsDNA-bound (**C**) states used in the molecular dynamics simulations. Trimeric systems with weak harmonic positional restraints on the backbone atoms of two outside protein chains and DNA were simulated. 100 snapshots with a time interval of 10 ns from 1 μs simulations are shown. The structures were fitted to the backbone atoms for the monomer in the middle of the trimeric system. Blue, silver and red colors correspond to early, mid and late time intervals, respectively.

As we hypothesized, the MD results clearly demonstrate the flexibility of this loop in all simulations. However, the greatest range of movement, as indicated by the highest average RMSD, was seen in the absence of DNA (Fig. 6A and Fig. S7 A, D-G). The conformations of the loop varied from completely open, which would allow the diffusion of the incoming DNA strand into the binding site, to completely closed, which would keep the DNA in there. The range of motion decreases considerably in the presence of ssDNA and further still, in the presence of nascent dsDNA (Fig. 6B-C, Fig. S7B-G). The decrease in motion is likely due to steric restrictions between the protein and the DNA. However, the flexible loop-tip contains two arginine residues, R134 and R137 (Fig. 5D), which may interact with the phosphate backbone of the incoming DNA strand to facilitate base pairing and thereafter helping to clamp the nascent dsDNA in place.

## Discussion

Despite its discovery in 1966 (*7*), the structure of Redβ annealase has remained unknown. It has proven to be extremely difficult to crystalize Redβ, despite trying a variety of truncations and added substrates. The crystals that could be grown did not diffract well enough to yield a structure (personal communication with Xinhua Ji at NIH and Charles Bell at OSU). Redβ_177_ used in this study was first described by the Bell group in 2006 (*49*). Our initial cryo-EM trials with apo protein yielded solenoid helices that were relaxed, and therefore, heterogeneous. Upon the addition of DNA and the optimization of buffer conditions, we have obtained compact solenoid helices of DNA-bound Redβ_177_ that are regular enough for helical image processing. The structure we have obtained unravels the molecular mechanisms of many observations and results published during the half a century of time since Redβ was discovered.

The first structural information of an annealase came from human Rad52 (*32, 45*). Redβ and Rad52 share only ∼17% amino acid sequence identity and previous bioinformatic studies placed the two proteins into superfamilies of their own (*3, 44*). However, similarity is evident at the level of the 3D structures (Fig. 3A, B) and the DALI server identified Rad52 as the most similar structure to Redβ_177_. Structures of Rad52 bound to ssDNA have been reported in two different modes, described as ssDNA bound to either the inner or outer DNA binding groove (*46*). Comparing our structure of DNA-bound Redβ_177_ to the Rad52 structures, we see that the placement of the template ssDNA in Redβ_177_ is similar to the ssDNA found at the inner binding site of Rad52. Both proteins have DNA-binding sites made up of the β-β-α-β-β-β-α secondary structure motif that is predicted to be conserved among the annealases (*43*). However, a major difference between the two proteins is the location of the flexible loop that is proposed to hold the DNA in place. In Redβ, the flexible loop forms the ‘top’ of the DNA binding site, whilst in Rad52 a functionally similar loop is found at the ‘bottom’ of the DNA binding site, located between the first two β-strands in the motif (Fig. 3A, B). The disparity between sequence and structure similarity between Redβ and Rad52 could be explained by two scenarios: 1) Both proteins are homologous and share a common ancestor but have diverged significantly over time. However, as structure is more conserved than sequence (*57*), the proteins remained structurally similar despite divergent sequence evolution. 2) The proteins are not homologues, but their structure is biologically favorable for the role of an annealase. In this scenario, the similarities in protein structure would be the result of convergent evolution towards an enzymatic optimum.

DdrB from *Deinococcus radiodurans*, which was originally identified as belonging to the Rad52 superfamily of annealases (*3*), has also been captured as a DNA annealing intermediate structure. Two pentameric DdrB rings assemble to form a DNA-binding channel bringing complementary ssDNA strands together facing each other in the middle of the double-ring complex (*58*). Based on the DdrB structure, when we saw the conversion of the Redβ_177_ solenoid structure from relaxed to tight following the addition of complementary ssDNA strands, we predicted that ssDNAs would be bound in between the ‘rings’ of the helix (i.e., lock-washers forming each turn of the helical filament), with the DNA bases facing each other. We had attributed compaction of the solenoid to annealing between ssDNA strands, bringing the lock-washer rings together. Therefore, the location of the DNA in our reconstructed map, (wrapped around the outside of the helical Redβ_177_ filament) was an unexpected observation. A previous study proposed that ssDNA bound around the outside of Redβ rings but that annealed dsDNA was located on the inside of the helical filament (*25*). We show that the predicted location of the ssDNA was correct, and that this DNA-binding groove is also the location of the annealed dsDNA. What is similar between the DdrB and Redβ is the conformation of the DNA in the annealing intermediate: the nascent dsDNA is found in a planar conformation, unlike the various helical conformations dsDNA forms in solution (*56*). This also explains why Redβ does not bind to pre-annealed dsDNA (*24*), as the double-helical structure known as the B-form DNA would not fit into the DNA binding site of Redβ.

The homo-oligomerization of Redβ has been reported by several groups using various approaches and techniques, including native mass spectrometry (*27*), electron microscopy (*25, 59*) and atomic force microscopy (*11, 26*). These results are briefly summarized above and in more detail in our recent review (*16*). Our reconstruction shows how Redβ_177_ oligomerizes with substantial contacts between adjacent subunits in the helix, but there is limited top-bottom contacts between corresponding monomers in the next turn of the helix (Fig. 4). This may explain the heterogeneity in the compactness of the solenoid-like helices of Redβ_177_ observed in the micrographs. Our structure also shows how DNA binding reinforces interactions between adjacent monomers, favoring oligomerization (Fig. 5B). There are 4 bases per monomer in the DNA-binding groove, but each monomer makes contacts with the phosphates from 6 nucleotides of the template strand (Fig. 5A, B). This may explain the discrepancy in the literature regarding the site size of Redβ, which has been reported as 4-5 nts (*48, 60*).

Our structure has elucidated an extensive hydrogen bonding network involved in DNA binding; validating predictions made in previous studies. In particular, highly conserved R161 forms a hydrogen bond with position 4 of the template strand (Fig. 5B), supporting previous observations that mutation of this residue is correlated with a 1000-fold reduction in annealing efficiency *in vivo (43*). Lysines at positions 36, 61, 111, 132, 148, 154, and 172 were reported to be critical for DNA binding *in vitro* and recombination *in vivo*. K61 was specifically reported to be critical for DNA annealing but not for initial ssDNA binding, and a role in binding to the incoming strand of DNA was suggested (*61*). Our structure shows that lysines 36, 61, 132, and 154 are located close to the DNA binding site, but 111 is not (K172 is missing from our structure). In addition, K69, which was not indicated to be involved in DNA-binding previously, is also shown to interact with the DNA (Fig 5B). Interestingly, K61 is closer to the template DNA than to the incoming DNA strand, while lysines 36, 69, 132 are closer to the incoming strand.

Although K148 is within close proximity to the DNA binding site, its side-chain points away from the DNA. Moreover, other positively charged residues H146, H153, R128 and R149 are also near the DNA, with R128 located within hydrogen-bonding distance of both DNA strands, while W143 can make a hydrogen-bond with the template strand.

Redβ catalyzed annealing of homologous ssDNA was reported to be an apparent first order reaction (the half-time of the reaction was independent of the DNA concentration) and a synaptic mechanism was suggested in which the DNA molecules are first brought together irrespective of homology, which is then followed by homology searching and base pairing (*60*). The annealing intermediate structure of Redβ_177_ that we determined reveals the molecular mechanism for a first-order annealing reaction. Our sample was generated by the sequential addition of the two complementary 27 base pair oligonucleotides to Redβ_177_ and the structure presented suggests that the binding of the template strand to Redβ is required as the initial step. After this, we hypothesized that the following events may be taking place: the incoming complementary ssDNA can bind transiently while base-pairing (and therefore, homology) between the two DNA strands is being evaluated, possibly held in place by the flexible loop (composed of residues 130-139 most of which are missing in our structure) to which we refer to as the ‘finger-loop’.

Once the finger-loop is reopened, the incoming ssDNA would stay in place if a strong enough base-pairing is achieved; otherwise, it would diffuse out or slide to allow a new base-pair match to be tested. This opening-closing and 2D and/or 3D diffusion of the incoming ssDNA would be iterated until homologous pairing is established between the two ssDNA strands bound to Redβ.

To test our hypothesis for the action of the finger-loop, we ran molecular dynamics simulations. Our results showed that the mechanism we proposed is supported by the finger-loop moving between open and closed conformations. The motion of the finger-loop is likely a dynamic process that can be described as tapping behind the incoming ssDNA, keeping it in place to allow more time for the homology search to take place, but loose enough to allow sliding (2D diffusion) and thereby the detection of homology in situations where the nucleotide sequence may be shifted. The simulations also showed that Redβ is most flexible when it is not bound to any DNAs and then becomes more rigid when bound to the template ssDNA. Redβ is then further stabilized upon forming an annealing complex with two DNA strands. This is in agreement with the reports of Redβ forming stable DNA annealing complexes (*24*).

Redβ_177_ contains a 3_10_ helix between the finger loop and α5. 3_10_ helices are often located between an alpha helix and a flexible region, where the weaker i+3 h-bonding network is proposed to allow transient unravelling (*62*). Along with α5 this 3_10_ helix forms part of motif 6 (Fig. 3C and Fig. S5) which is highly conserved across annealases (Fig. S4). In addition, W143 of the 3_10_ helix is almost universally conserved across all species. Our structure identified that W143 forms part of the H-bonding network that stabilizes the induced kink in the phosphate backbone of the template DNA strand. Therefore, the position of W143 in this 3_10_ helix is likely to be of functional importance. The MD simulations indicated that the 3_10_ helix was unstable, oscillating through distorted conformations in concert with the movement of the finger-loop. Therefore, we propose a mechanism where transient relaxation of the 3_10_ helix as the finger-loop opens allows H-bonding of W143 to the phosphate backbone of the template strand. Then, as the loop closes and the 3_10_ helix reforms, the phosphate backbone may be pulled into the distorted conformation seen in our structure. The distortion in the phosphate backbone allows the template DNA strand to remain planar. This facilitates base pairing between the nucleotides of the template and incoming strands, both of which are held in a planar orientation through F66 (π-stacking) and R128 (H-bond), enhancing the formation of a stable annealing intermediate.

Finally, we have created a molecular scene by piecing four structures together — 4WUZ (λExo with DNA (*63*)), 6M9K (λExo with the C-terminal domain of Redβ (*64*)), our Redβ_177_ model, and the AlphaFold prediction (Redβ_AF_ (*51*)) of full-length Redβ (Fig. S8). This scene (Fig. 7) shows dsDNA bound to λExo, from which the digested nascent ssDNA would be extruded from the center of the λExo toroid towards the Redβ molecules bound to the ‘back’ side of λExo (only one Redβ is shown, but there would be 3, one for each λExo monomer). The Redβ_AF_ prediction shows helix α7 placed near the DNA-binding pocket of Redβ. Therefore, the superposition of the Redβ_177_ and Redβ_AF_ places α7 in a position that is partially clashing with the bound DNA. If AlphaFold is correctly predicting the position of this helix in the apo state, our structure of Redβ shows that helix α7 must be displaced via a conformational change to allow DNA binding. This may function as an auto-inhibition mechanism, to ensure that Redβ would not bind to ssDNAs generated in the cell during other processes, such as DNA replication, and that it is only able to bind to nascent ssDNA generated via digestion of dsDNA by λExo. The nascent ssDNA would be effectively “loaded” onto Redβ by this coupling mechanism, coordinating the two reactions catalyzed by the individual components of this two-component recombinase: DNA digestion by λExo and DNA binding and annealing by Redβ. Our molecular scene provides a glimpse of what this coupling may look like, furthering our understanding of the molecular mechanisms of SSA.

**Fig. 7.**
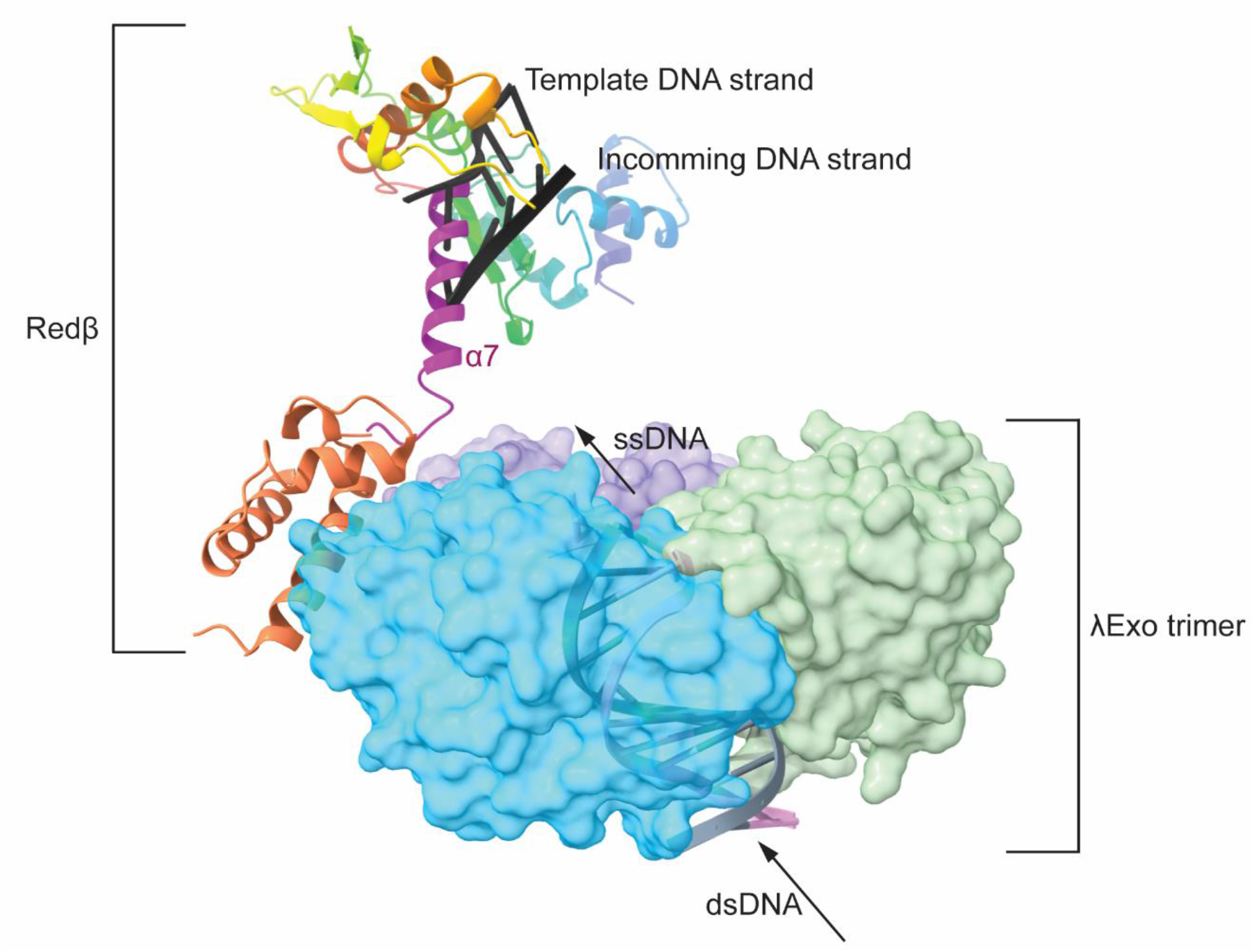
Superimposed molecular structure scene of Redβ, λExo and DNA. This molecular scene was created by piecing four structures together. These include the structure of λExo with DNA (*63*) (4WUZ), the structure of λExo with C-terminal domain of Redβ (*64*) (6M9K), the Redβ_177_ model and the AlphaFold prediction for the full length Redβ (*51*). Redβ is shown as a ribbon structure, while λExo is represented as a surface render.

There are many questions that remain about the molecular mechanisms of SSA, some as mysterious as “How can a trimeric λExo (3 subunits) and a dodecameric Redβ (12 subunits) form a complex with 1:1 monomer:monomer stoichiometry?” or “How does Redβ form rings and helices on ssDNA in concert with dsDNA digestion by λExo?”.

Moreover, the structure we have determined poses new questions, such as “what is the function and significance of the finger-loop in Redβ and Rad52?” and “how is nascent dsDNA released from stable Redβ helices back into solution?”. As efforts to answer the remaining questions are still ongoing more than half a century after its discovery, our structure of Redβ_177_ finally provides insights into the mechanism of how this protein functions.

## Materials and Methods

### Cloning of Redβ_177_

A pET-28a(+) plasmid containing the wild-type *bet* gene was kindly provided by the laboratory of Dr. Donald Court (National Cancer Institute, NIH, Frederick, MD, USA). Redβ_177_ was assembled into a pET-24a(+) plasmid for expression with a C-terminal hexa-histidine (His_6_) tag. The DNA sequence corresponding to Redβ_177_ was PCR amplified using primers 5’-GTTTAACTTTAAGAAGGAGATATACCATGAGTACTGCACTCGCAAC-3’ and 5’-GATCTCAGTGGTGGTGGTGGTGGTGAGAAGAGCGCTCGGCTTCATCCTTG-3’, for the simultaneous addition of pET-24a(+) overlap sequences. The pET-24a(+) plasmid was PCR amplified using primers 5’-GGTATATCTCCTTCTTAAAGTTAAACAAAATTATTTCTAG-3’ and 5’-TCTTCTCACCACCACCACCACCACTGAGATC-3’. Following PCR, both DNA fragments were purified by agarose gel electrophoresis and subsequent extraction. The recombinant plasmid was generated by a NEBuilder^®^ HiFi DNA Assembly (New England Biolabs) reaction, under standard conditions and using a 2:1 molar ratio of insert:plasmid DNA. The resulting construct, pET24a-bet_177_-His_6_, places the expression of Redβ_177_-His_6_ (Redβ_177_) under the control of the *E. coli lac* expression system. The correct sequence of the plasmid was confirmed by Sanger sequencing (Garvan Molecular Genetics, Garvan Institute of Medical Research, Sydney, Australia). *E. coli* strain BL21(DE3) was transformed by the recombinant plasmid and used for protein expression.

### Expression and purification of Redβ_177_ protein

*E. coli* BL21(DE3) cells were cultured in LB media supplemented with 50 μg/ml kanamycin, at 37°C and with shaking at 200 rpm. When the culture reached an OD_600 nm_ of 0.6, protein expression was induced by the addition of 1 mM isopropyl-β-D-1-thiogalactopyranoside (IPTG, Astral Scientific). At 3 hours post-induction, the cells were harvested by centrifugation at 6,100 *g* for 7 minutes at 6°C. The cell pellet was resuspended in 50 ml of Lysis buffer (50 mM Tris pH 7.5, 300 mM NaCl, 10% *v*/*v* glycerol, 10% *w*/*v* sucrose) and cells were lysed in an ice bath by sonic pulses produced by an Ultrasonics digital sonifier (Branson). The lysate was clarified by centrifugation at 38,000 *g* for 30 minutes at 4°C. Following centrifugation, the supernatant was passed through a 0.45 μm syringe-driven filter (Sarstedt).

At a flow rate of 1 ml/min, controlled by an ÄKTA Pure FPLC equipped with a sample pump (Cytiva), the filtrate was pumped through a 1 ml HisTrap HP column (Cytiva), pre-equilibrated with 50 mM Tris pH 7.5, 300 mM NaCl, 10% *v*/*v* glycerol and 10 mM imidazole. The flow-through from the column, containing unbound proteins, was collected and retained. The column was washed at 1 ml/min with 10 column volumes of wash buffer (50 mM Tris pH 7.5, 300 mM NaCl, 10% *v*/*v* glycerol and 20 mM imidazole), before bound proteins were eluted with a 0-100% gradient of elution buffer (50 mM Tris pH 7.5, 300 mM NaCl, 10% *v*/*v* glycerol and 500 mM imidazole), over 30 column volumes at a flow rate of 1 ml/min. The column was then washed with 10 column volumes of elution buffer, before being re-equilibrated.

The flow-through from the first separation was pumped back onto the column for further extraction of Redβ_177_. Elution fractions from both chromatography separations were analyzed by SDS-PAGE. Fractions that were highly enriched for Redβ_177_ were pooled and concentrated in a 10 kDa MWCO centrifugal concentrator (Merck Millipore) to a total volume of <2 ml. The concentrated protein was centrifuged at 21,000 *g*, for 30 min at 4°C, following which there was no visible pellet. The supernatant was injected onto a HiLoad 16/600 Superdex pg gel filtration column (Cytiva), equilibrated with 50 mM Tris pH 7.5, 50 mM NaCl, 10% *v*/*v* Glycerol. Redβ_177_ eluted as a single, symmetric peak at 1 ml/min flow. The central peak fractions were pooled and estimated to be <95% pure by SDS-PAGE. Protein concentration was assessed by measuring absorbance at 280 nm using a NanoDrop 2000c spectrophotometer (ThermoFisher Scientific) and the extinction coefficient of ε = 34950 L.mol^-1^.cm^-1^. Protein was stored stable for up to 3 weeks at 4°C, or flash frozen in liquid nitrogen and then stored at -80°C.

### Helical complex assembly and preparation of cryo-EM grids

To form the helical assemblies, purified Redβ_177_ was incubated with complementary 27mer oligonucleotides. Redβ_177_ was incubated with Oligo 1 (5’-TGCAGCAGCTTTACCATCTGCCGCTGG-3’) in binding buffer (20 mM KH_2_PO_4_ pH 6.0, 5 mM MgCl_2_) for 30 minutes at room temperature. Oligo 2 (5’-CCAGCGGCAGATGGTAAAGCTGCTGCA-3’) was then added, and the incubation was continued for a further 30 minutes. A molar ratio of 7.7:1:1 of Redβ_177_:Oligo 1:Oligo 2 was used. To remove any remaining traces of glycerol from the storage buffer, the complex was dialyzed against the binding buffer overnight at 4°C, using a 10 kDa MWCO Slide-A-Lyzer MINI Dialysis cassette (ThermoFisher Scientific). A 2.5 μL aliquot of the dialyzed complex was deposited onto gold UltrAuFoil R1.2/1.3 Au 300 mesh cryo-electron microscopy grids (Quantifoil) and plunge frozen into liquid-nitrogen cooled liquid ethane using a Mark IV Vitrobot (ThermoFisher Scientific).

### Cryo-electron microscopy and helical reconstruction of Redβ_177_-6xHis helical assemblies

The data was collected using a Titan Krios electron microscope (ThermoFisher scientific) which operated at 300 kV. The Titan Krios was equipped with a Gatan K2 Summit detector and a Gatan BioQuantum LS 967 energy filter, operated in unfiltered mode. Data was collected in electron counting mode, using a pixel size of 0.84 Å/pixel as well as a calibrated sample-to-pixel magnification of 59524 x.

Movies were collected as a series of 50 frames with a total accumulated dose of 50 e^−^/Å^2^. Data were collected using automated data collection in EPU, with a defocus range of -0.6 to -2.2 μm. All processing was performed in cryoSPARC (*52*) unless otherwise noted. Movies were gain corrected and aligned using patch-based motion correction and patch-based CTF correction was performed. Helical filament tracing and extraction was performed with a box size of 576 pixels and a particle separation distance of 55 pixels. Extracted particles were subjected to 2D classification and subsequent 2D class averages were used as templates for further rounds of template-based helical filament tracing. This process was repeated twice until picking had reached sufficient coverage of the helical filaments. Particles were then re-extracted with a box size of 448 pixels and rescaled to a box size of 224 pixels to assist with the speed of reconstruction. Extracted particles were subjected to several rounds of 2D classification to identify suitable subsets for reconstruction.

Suitable classes identified via 2D classification were initially subjected to 3D helical reconstruction with no helical symmetry applied. Once helical reconstruction had converged, particles were re-extracted to a box size of 448 pixels with no rescaling. Helical reconstruction was then repeated, and the helical symmetry parameters were estimated using real-space helical symmetry estimation. Helical reconstruction was then performed with helical symmetry and subsequent non-uniform refinement. CTF refinement and local motion correction were then performed until convergence. The final resolution of the reconstructed density was estimated using the gold-standard Fourier shell correlation (GS-FSC) criteria.

### Structure building and refinement

*De novo* atomic modelling in COOT (*65*) was used to generate the initial structure for the Redβ_177_ monomer and the bound nucleic acid. The model generated in COOT was then refined using ISOLDE (*66*). The atypical conformation of the template and non-template strands was modelled with the assistance of adaptive distance restraints in ISOLDE. Model statistics were assessed via MolProbity (*67*) as implemented in the Phenix software suite (*68*).

### Bioinformatics

In order to thoroughly search the available sequence data for homologous sequences, a full length Redβ amino acid sequence (UniProtKB: P03698) was BLAST searched (E-value cut-off = 1, substitution matrix = BLOSUM-64) against the UniProt Reference Cluster (UniRef) 90% database (*69*). From this search, the top 1000 hits were extracted, and amino acid sequences aligned with the command line version of MAFFT (**<u>M</u>**ultiple **<u>A</u>**lignment using **<u>F</u>**ast **<u>F</u>**ourier **<u>T</u>**ransform) v.7.480 (*70*) using the L-INSI algorithm. All MAFFT parameters were left at default, except for the ‘maxiterate’ parameter that was set to 1000.

The resultant multiple sequence alignment (MSA) was then visualized within Jalview v.2.11.1.4 (*71*) and trimmed to the region of Redβ containing the residues 19 – 205. Redundant sequences and sequences less than 100 residues were then removed. Outlier sequences were also identified using the principal component and neighbor joining tree analysis tools within Jalview (both using the default settings) and removed. Following refinement, the MSA was then again aligned with MAFFT using the above parameters. Conserved motifs were then identified based on 50% consensus across the MSA as calculated in Geneious Prime v.2021.1.1 (available from: https://www.geneious.com/). Conserved motifs were then manually refined by identifying, and subsequently annotating, positions at which an amino acid property, rather than an individual residue, was conserved across the MSA.

DALI protein structure comparison server (*55*) was used with the default parameters for identifying the proteins that are structurally similar to Redβ_177_.

### Molecular dynamics simulations and structure prediction

Molecular dynamics simulations were carried out using NAMD (**<u>Na</u>**noscale **<u>M</u>**olecular **<u>D</u>**ynamics, version 3 alpha) (*72*). In total, three systems have been simulated each for 1 μs, including Redβ alone (apo), Redβ bound to ssDNA (ss), and Redβ bound to the dsDNA annealing intermediate (ds). For each system, two different setups have been simulated: one set of the simulations are free simulations, while the other set of simulations are applied a positional harmonic restraint on the two outside protein and DNA backbone atoms. The latter was used to probe the effects of using trimeric structure to study the oligomeric structure. Initial preparation of systems was done with CHARMM-GUI (*73, 74*). The missing loops (residue 131 to 136) were built with Modeller (version 10.1) (*75*). The AMBER protein parm14SB force field (*76*) and nucleic acid BSC1 force field (*77*) were applied for the protein and DNA respectively. The TIP3P model was used for water (*78*).

Molecular dynamics simulations were performed after solvating the system in an ∼100×100×100 Å cubic box that extended at least 10 Å from the solute surface. Na^+^ and Cl^−^ counter ions were added to neutralize the system and achieve a salt concentration of 0.15 M. pKa, calculations were performed using PROPKA to assign protonation states of ionizable residues. Simulations were performed using periodic boundary conditions (PBC) at constant temperature (303.15 K) with the Langevin algorithm (a damping coefficient of 1/ps) (*79*) and at a pressure of 1.0 bar using the Nose-Hoover Langevin Piston method (*80*). The time step was set to 2.0 fs with all covalent bonds involving hydrogens kept rigid with the RATTLE algorithm (*81*). Short-range electrostatics were calculated together with long-range electrostatics particle mesh Ewald (PME) (*82*) with a cutoff of 9.0 Å and a PME grid size of 1.0 Å. For all systems, energy minimization (10,000 steps) and 125 ps equilibration were performed first with positional restraints placed on all the protein and DNA heavy atoms (with a force constant of 1.0 kcal/mol/Å^2^ on the backbone atoms and 0.5 kcal/mol/Å^2^ on the side chain atoms). This was followed by 1 μs production runs. In the simulation set with a positional harmonic restraint, this restraint was maintained on the two outside monomers and DNA backbones. Snapshots were saved every 100 ps. VMD (**<u>V</u>**isual **<u>M</u>**olecular **<u>D</u>**ynamics) (*83*), LOOS (**<u>L</u>**ightweight **<u>O</u>**bject-**<u>O</u>**rientated **<u>S</u>**tructure Analysis) (*84*) and MDAnalysis (*85*) were used to analyze the trajectories.

Structural prediction was performed with the ColabFold (*86*) implementation of AlphaFold2. The source code was retrieved from the GitHub repository (https://github.com/YoshitakaMo/localcolabfold). For each prediction, five models were generated and optimized with the AMBER force field.

## Acknowledgements

We would like to thank Dr. James Bouwer and Dr. Simon Brown at the Molecular Horizons Cryo-Electron Microscopy Facility, and the MASSIVE (Multi-modal Australian ScienceS Imaging and Visualisation Environment) team for their technical help and support during image data collection and processing. Dr. Tolun extends his thanks to Dr. Donald Court for his support during the early stages of this work.

