## Supplementary figures for "Structure of phage λ Redβ_177_ annealase shows how it anneals DNA strands during single-strand annealing homologous DNA recombination"

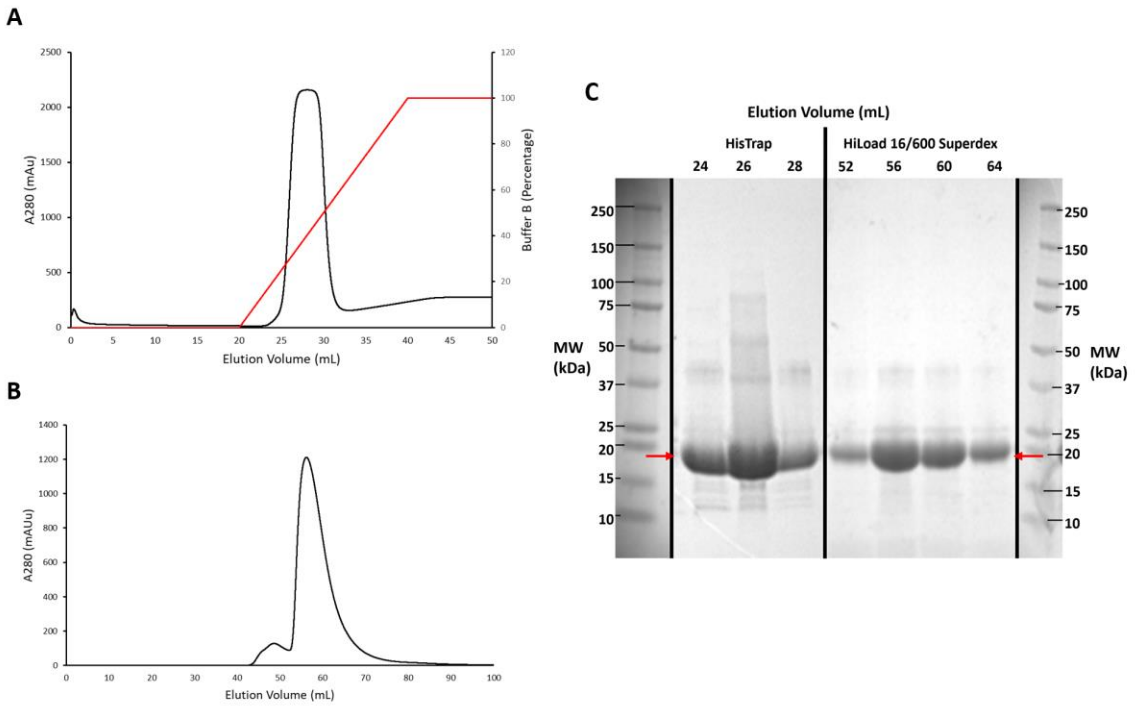


**Fig. S1. The two step purification of Redβ$_{177}$.** (**A**) Column profile for immobilized metal ion affinity chromatography of Redβ$_{177}$. (**B**) Column profile for size exclusion chromatography purification through a HiLoad 16/600 Superdex column. The black line represents the UV (280 nm) absorbance, while the red line represents the increasing the concentration (%) from 10 mM to 500 mM imidazole. (**C**) SDS-PAGE gel showing bands at the expected size of Redβ$_{177}$, highlighted by red arrows.

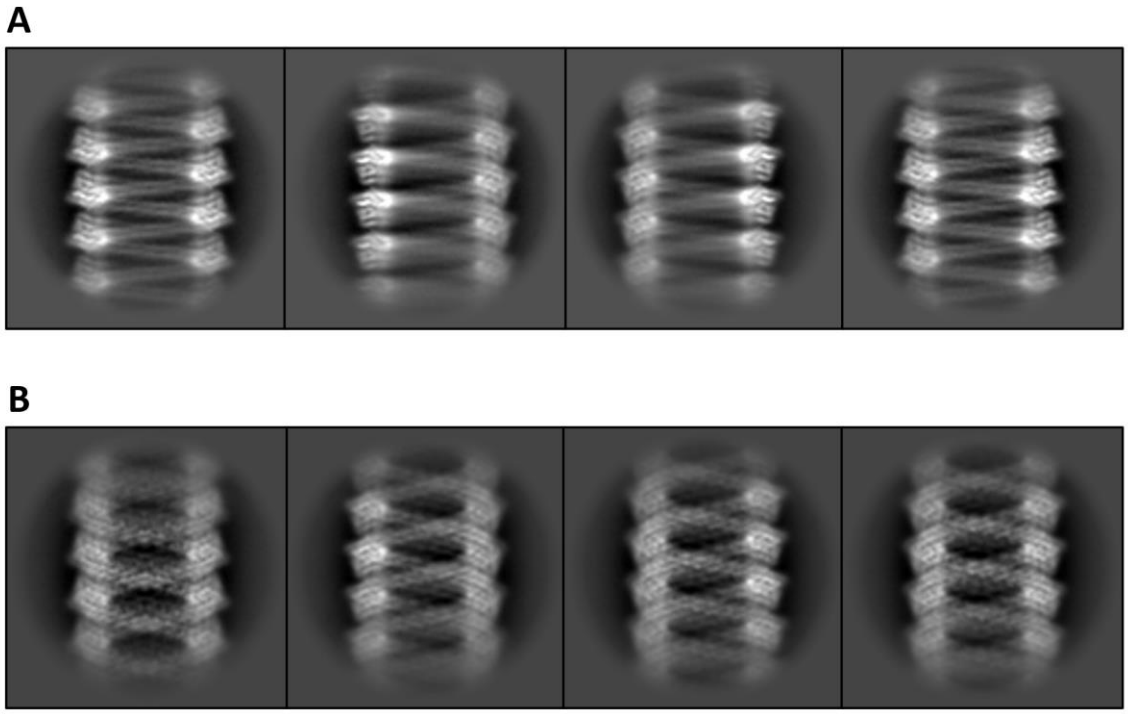

**Fig. S2. 2D class averages for the 1-start and 2-start helical assemblies.** Selected 2D class averages of the Redβ$_{177}$ helical filaments obtained after processing in cryoSPARC (*1*). These include populations of both the 1-start helix (**A**) and the 2-start helix (**B**).

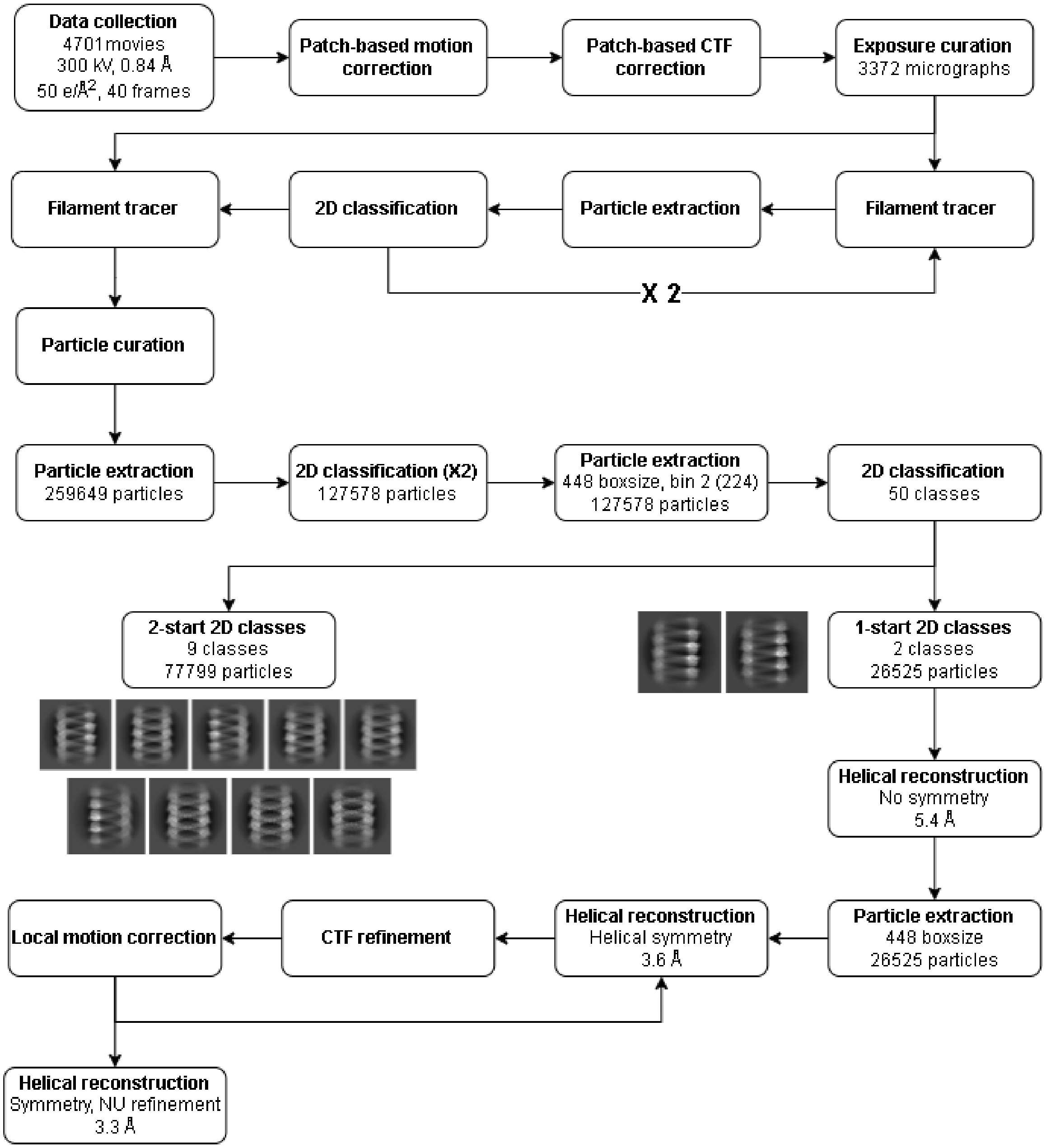


**Fig. S3. Overall workflow of the cryo-EM data processing.** Flowchart shows the cryo-EM data analysis performed, from the initial micrographs to the final reconstruction.

**Fig. S4. Multiple Amino Acid Sequence Alignment (Data file uploaded).** The alignment is composed of the top 1000 UniProt Ref90 clusters related to the full-length Redβ amino acid sequence (UniProtKB: P03698). Conserved sequence motifs, based on 50% consensus, are shown (top) with individual residues colored by amino acid side chain property: hydrophobic = orange, negative charge = red, polar uncharged = pink, positive charge = blue and other properties/ambiguous amino acid codes = black. Positions where an amino acid property is conserved, rather than an individual residue, are indicated: hydrophobic = 'h', polar uncharged = 'p', positive charge = '+'. The eight conserved motifs (M1-M8) identified are indicated (top). The alignment is colored following the 'ClustalX' scheme as implemented in Jalview v.2.11.1.4 (*2*).

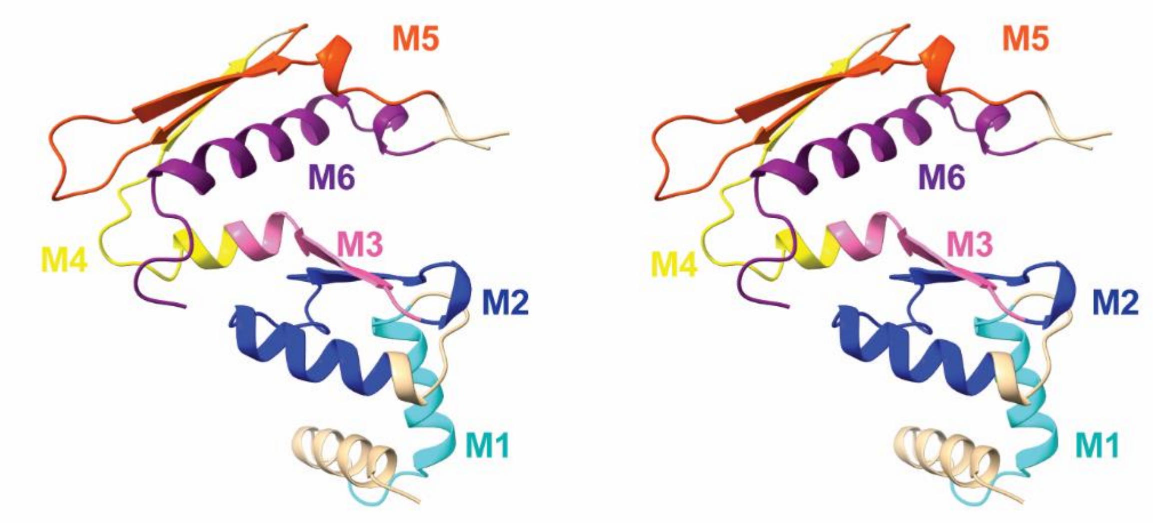


**Fig. S5. Wall-eye (parallel) stereo of the conserved motifs from the MSA mapped onto the structure of Redβ$_{177}$.** Colors indicate different conserved motifs identified by the MSA (Fig. S4), with 6 out of the 8 motifs being represented in separate colors and numbered. Beige indicates the areas outside of the identified motifs.

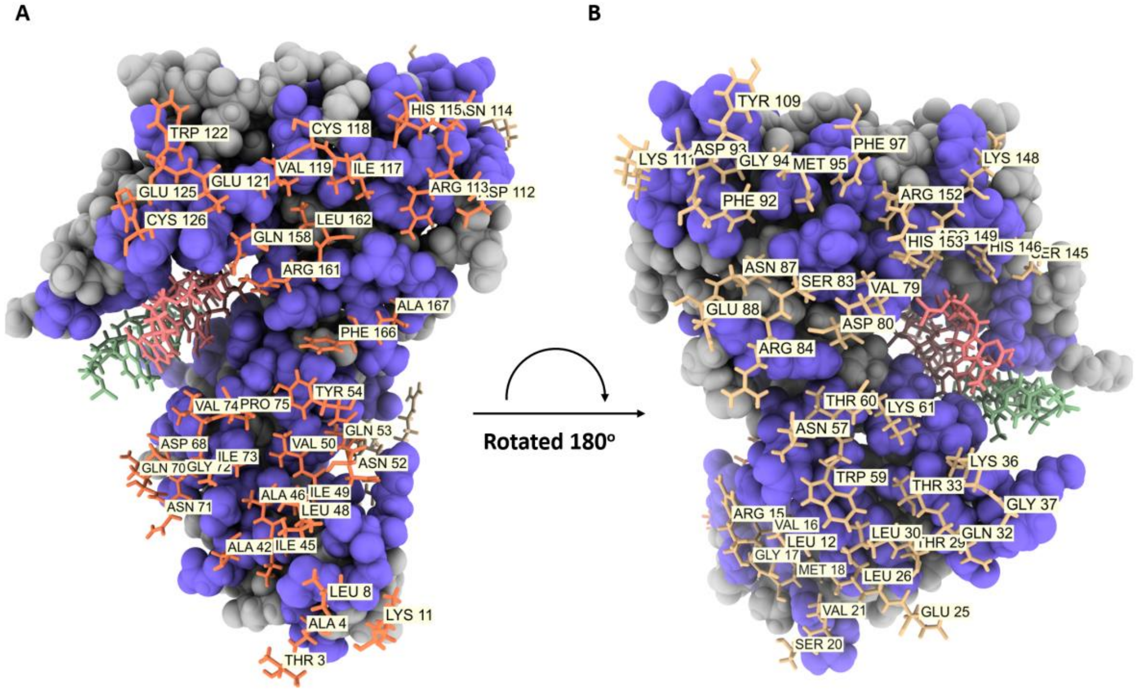


**Fig. S6. Redβ177 Interacting residues.** Residues involved in side-by-side interactions between two Redβ177 monomers. (**A**) and (**B**) show the two sides of the Redβ177 monomer that would be facing each other in the helical assembly.

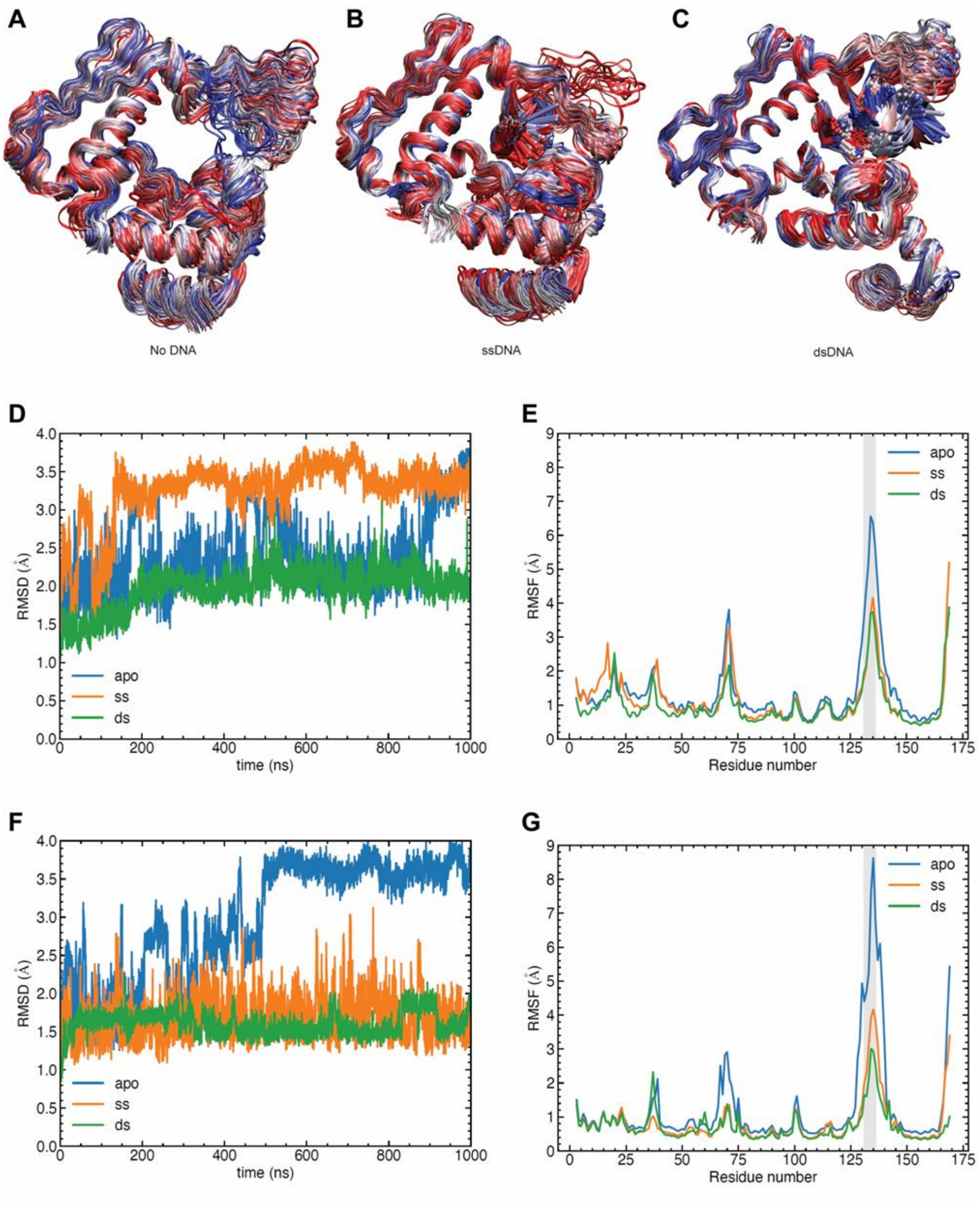


**Fig. S7. Molecular Dynamics without restraints snapshots.** The sampled ensemble of apo (**A**), ssDNA-bound (**B**) and dsDNA-bound (**C**) $\mathrm{Red}\beta_{177}$ states in the MD simulations of the trimeric systems without any restraints. 100 snapshots with a time interval of 10 ns from 1 μs simulations are shown. The structures were fitted to the backbone atoms for the monomer in the middle of the trimeric system. Blue, silver and red colors correspond to early, mid and late time intervals, respectively. The backbone positional root-mean-square deviations for the middle protein chain from the initial structure as a function of simulation time (**D**) without any restraints and (**F**) with a weak harmonic restraints on the two outside monomer and DNA backbone atoms). The Cα atom positional root-mean-square fluctuations for the middle protein chain (**E**) without any restraints and (**G**) with a weak harmonic restraints.

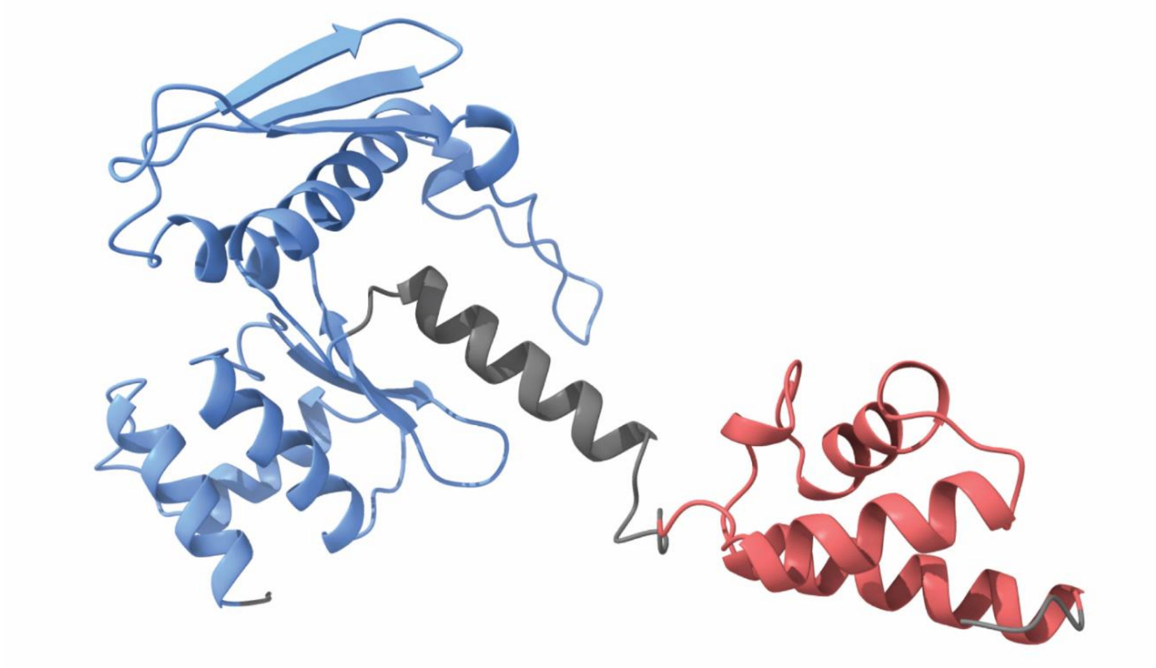

**Fig. S8. The full-length Redβ structure predicted by AlphaFold** (*3*). Blue shows the N-terminal domain that corresponds to the Redβ$_{177}$ structure in this study, red shows the residues that correspond to the C-terminal domain of Redβ determined by Caldwell *et al.* (*4*), and grey are the residues whose structures were not experimentally determined.

**Table S1. Cryo-EM data collection, refinement and validation statistics.**

| | Redβ177-dsDNA (PDB: 7UJL, EMD-26566) |
|---|---|
| Data collection and processing | |
| **Molecular weight (kDa/nm)** | 114.18 |
| **Magnification (x)** | 59500 |
| **Voltage (kV)** | 300 |
| **Electron exposure (e-/Å2)** | 50.0 |
| **Defocus range (Å)** | 500-2500 |
| **Pixel size (Å/pixel)** | 0.84 |
| **Initial particle images** | 259,649 |
| **Final particle images** | 26,525 |
| **Axial symmetry imposed** | C1 |
| **Helical rise (Å)** | 2.078 |
| **Helical twist (°)** | -12.947 |
| **Map resolution (Å)** | 3.1 |
| **FSC threshold** | 0.143 |
| **Map resolution range (Å)** | 2.8-4.8 |
| Refinement | |
| **Initial model used** | N/A |
| **Model resolution (Å)** | 3.2 |
| **FSC threshold** | 0.5 |
| Model composition | |
| **Non-hydrogen atoms** | 1447 |
| **Protein residues** | 161 |
| **Nucleic acid residues** | 8 |
| B factors | |
| **Protein** | 30.00 |
| **Nucleic** | 9.27 |
| r.m.s deviations | |
| **Bond lengths (Å)** | 0.015 |
| **Bond angles (°)** | 2.075 |
| Validation | |
| **MolProbity score** | 0.85 |
| **Clashscore** | 0.00 |
| **Poor rotamers (%)** | 0.00 |
| Ramachandran plot | |
| **Favoured (%)** | 94.90 |
| **Allowed (%)** | 5.10 |
| **Outliers (%)** | 0.00 |

**Supplementary Information References**


1. A. Punjani, J. L. Rubinstein, D. J. Fleet, M. A. Brubaker, cryoSPARC: algorithms for rapid unsupervised cryo-EM structure determination. *Nat Methods* **14**, 290-296 (2017).
2. A. M. Waterhouse, J. B. Procter, D. M. Martin, M. Clamp, G. J. Barton, Jalview Version 2--a multiple sequence alignment editor and analysis workbench. *Bioinformatics* **25**, 1189-1191 (2009).
3. J. Jumper *et al.*, Highly accurate protein structure prediction with AlphaFold. *Nature* **596**, 583-589 (2021).
4. B. J. Caldwell *et al.*, Crystal structure of the Redbeta C-terminal domain in complex with lambda Exonuclease reveals an unexpected homology with lambda Orf and an interaction with Escherichia coli single stranded DNA binding protein. *Nucleic Acids Res* **47**, 1950-1963 (2019).
