## Supplementary figures and images for "Structure of phage λ Redβ_177_ annealase shows how it anneals DNA strands during single-strand annealing homologous DNA recombination"

### Supplementary figure 4

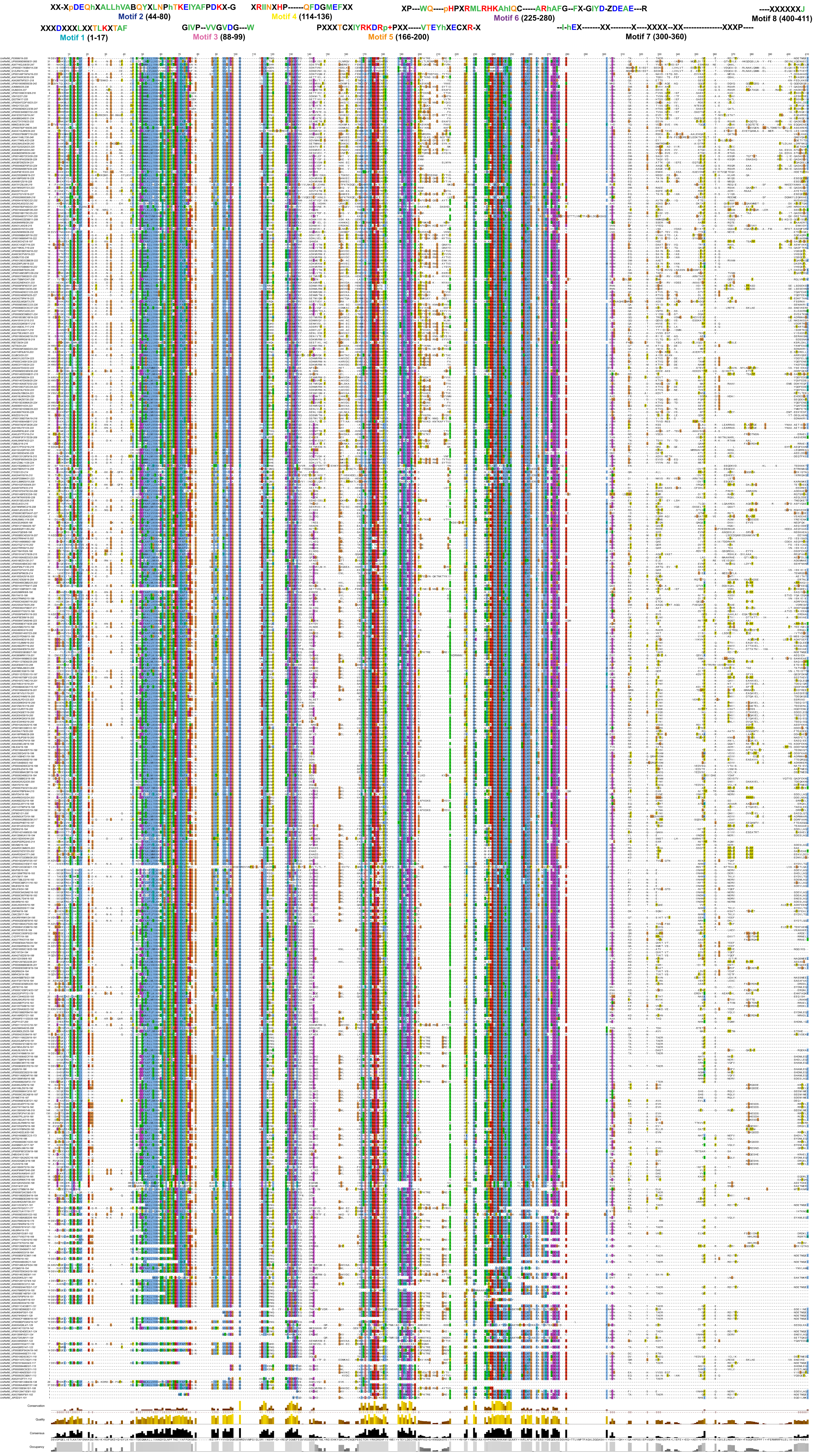
